# CD4 T cells are rapidly depleted from tuberculosis granulomas following acute SIV co-infection

**DOI:** 10.1101/2021.12.17.473203

**Authors:** Taylor W. Foreman, Christine E. Nelson, Keith D. Kauffman, Nickiana E. Lora, Caian L. Vinhaes, Danielle E. Dorosky, Shunsuke Sakai, Felipe Gomez, Joel D. Fleegle, Melanie Parham, Shehan R. Perera, Cecilia S. Lindestam Arlehamn, Alessandro Sette, Tuberculosis Imaging Program, Jason M. Brenchley, Artur T.L. Queiroz, Bruno B. Andrade, Juraj Kabat, Laura E. Via, Daniel L. Barber

**Affiliations:** T lymphocyte Biology Section, Laboratory of Parasitic Diseases, National Institutes of Allergy and Infectious Disease, National Institutes of Health, Bethesda, MD, USA; Multinational Organization Network Sponsoring Translational and Epidemiological Research (MONSTER) Initiative, Instituto Gonçalo Moniz, Fundção Oswaldo Cruz, Salvador, Brazil; Bahiana School of Medicine and Public Health (EBMSP), Salvador, Brazil; Division of Intramural Research, National Institutes of Allergy and Infectious Disease, National Institutes of Health, Bethesda, MD, USA; Axle Informatics, National Center for Advancing Translational Sciences, Bethesda, MD, USA; Department of Electrical and Computer Engineering, The Ohio State University, Columbus, OH, USA; Division of Vaccine Discovery, La Jolla Institute for Immunology, La Jolla, CA, USA; Barrier Immunity Section, Laboratory of Viral Diseases, National Institutes of Allergy and Infectious Disease, National Institutes of Health, Bethesda, MD, USA; Data and Knowledge Integration Center for Health (CIDACS), Instituto Gonçalo Moniz, Salvador, Bahia, Brazil; Biological Imaging Section, Research Technologies Branch, National Institutes of Allergy and Infectious Disease, National Institutes of Health, Bethesda, MD, USA; Tuberculosis Research Section, Laboratory of Clinical Immunology and Microbiology, National Institutes of Allergy and Infectious Disease, National Institutes of Health, Bethesda, MD, USA; Institute of Infectious Disease & Molecular Medicine and Division of Immunology, Department of Pathology, University of Cape Town, Observatory, South Africa

**Author notes:** Note: The members of the NIAID/DIR Tuberculosis Imaging Program can be found at the end of the Acknowledgments. Corresponding author: Dr. Daniel Barber. Author Contributions: T.W.F. conceived and led the study. T.W.F. and D.L.B. designed the research. T.W.F., C.E.N., K.D.K., N.E.L., D.E.D, and S.S. performed the experiments. T.W.F., C. E.N., A.T.L.Q., C.L., B.B.A., and D.L.B. analyzed the Luminex results. T.W.F., C.E.N., and D. L.B. analyzed the flow cytometry results. T.W.F., J.K., and D.L.B. analyzed the 4D imaging results. M.P. and S.R.P. wrote scripts necessary for 4D imaging analysis. C.S.L.A., A.S., and J.M.B. provided critical reagents. Tuberculosis Imaging Program (TBIP) managed logistics, performed NHP manipulations including infection, necropsy, PET/CT scanning, and imaging analysis. F.G. and J.D.F. analysized the PET/CT data. L.E.V. supervised TBIP and designed the analysis for PET/CT data. T.W.F. and D.L.B. wrote the manuscript with all authors contributing with feedback.

## Abstract

The HIV-mediated decline in circulating CD4 T cells correlates with increased risk of active tuberculosis (TB)^1–4^. However, HIV/*Mycobacterium tuberculosis* (Mtb) co-infected individuals also have an increased incidence of TB prior to loss of CD4 T cells in blood^3,5^, raising the possibility that HIV co-infection leads to disruption of CD4 T cell responses at the site of lung infection before they are observed systemically. Here we used a rhesus macaque model of SIV/Mtb co-infection to study the early effects of acute SIV infection on CD4 T cells in pulmonary Mtb granulomas. Two weeks after SIV co-infection CD4 T cells were dramatically depleted from granulomas, before significant bacterial outgrowth, disease reactivation as measured by PET-CT imaging, or CD4 T cell loss in blood, airways, and lymph nodes. Mtb-specific CD4 T cells, CCR5-expressing, in granulomas were preferentially depleted by SIV infection. Moreover, CD4 T cells were preferentially depleted from the granuloma core and lymphocyte cuff relative to B cell-rich regions, and live imaging of granuloma explants showed that SIV co-infection reduced T cell motility. Thus, Mtb-specific CD4 T cells in pulmonary granulomas may be decimated before many patients even experience the first symptoms of acute HIV infection.

## MAIN

Human immunodeficiency virus (HIV) infection gradually depletes circulating CD4 T cells leading to the development of acquired immunodeficiency syndrome (AIDS) and impaired host resistance to microbial infections. CD4 T cells are essential for control of *Mycobacterium tuberculosis* (Mtb) infection, and tuberculosis (TB) is the leading cause of death in persons living with HIV (PLWH)^6^. The extent of peripheral CD4 T cell depletion in PLWH correlates with increasing risk of developing active TB^3,4^. However, PLWH with normal-to-high CD4 T cell counts in the blood also have increased incidence of active TB^3^. The mechanistic basis for the elevated risk of TB in individuals with normal CD4 T cell counts is incompletely understood, but may reflect detrimental impact of antiviral inflammation (e.g. type I interferons) on anti-mycobacterial innate immunity^7–9^, impairment of macrophage function^10,11^, or preferential depletion of CD4 T cells in tissues compared to circulation^12–14^.

Mtb persists in granulomas, complex structures comprised of multiple immune cell types that reproducibly position themselves relative to one another^15^. Although variable by many respects, granulomas can be generally described by a few key features including: an of-tnecrotic, macrophage-rich, core where mycobacteria are primarily located^16^, a lymphocyte-rich cuff circumscribing the macrophage core^17^, and B cell-rich tertiary lymphoid-like substructures referred to as Granuloma-Associated Lymphoid Tissue (GrALT) located within the lymphocyte cuff or found as proximal granuloma satellites^18,19^. CD4 T cells are primarily found in the lymphocyte cuff but can also be seen interacting with macrophages in the core or with B cells in GrALT^17^. Due to the difficulty of studying human lung infections, little is understood about the effects of HIV infection on Mtb granulomas. Previous studies have shown that SIV infection of macaques recapitulates the gradual loss of circulating CD4 T cells and eventual development of active tuberculosis disease as seen in humans^20–23^. Here we examine the very early effects of SIV infection on CD4 T cell responses at the site of Mtb infection in rhesus macaques to ask if CD4 T cells within the granuloma microenvironment are differentially susceptible to dysfunction and depletion during SIV infection as compared to T cells in circulation or lymphoid tissues.

## RESULTS

To study the early effects of SIV co-infection in Mtb-infection, 10 Indian rhesus macaques were intrabronchially infected with ~56 CFU of *Mtb* H37Rv bilaterally (Fig. 1a). Animals were monitored for signs of disease for 10-11 weeks before allocation into groups that either remained Mtb *mono*-infected (n=5) or were intravenously infected with 3,000 TCID_50_ of SIVmac239 (n=5). The study was concluded exactly fourteen days after SIV infection. Animals did not show signs of tuberculosis disease as measured by weight loss or fever over the course of the study (Fig. S1a-b). Plasma SIV viral loads increased in most animals until day 14 (Fig. 1b). Using RNAscope to study SIV localization, viral RNA was detected in granulomas, surrounding lung tissue, and lymph nodes (Fig. 1c-d). (Fig. 1d, S2). Spot density of RNAScope for SIV RNA was higher in granulomas than lymph node and surrounding lung tissues (Fig. 1e). Spots of viral RNA staining in granulomas were further analyzed for their distribution in the core, cuff, or GrALT (confirmed as CD20-rich areas). Viral RNA staining was most dense in GrALT regions as compared to the core and cuff of the granuloma (Fig. 1f). Although it is difficult to distinguish between viral particles and infected cells with this technique, we did find that the size of individual spots was significantly different between tissues. Lymph nodes contained a higher frequency of large spots >100 μm^2^ (possibly representing infected cells) compared to the granulomas (Fig. 1g). Overall, these results demonstrate that granulomas become heavily infected with virus quickly after SIV infection.

**Figure 1.**
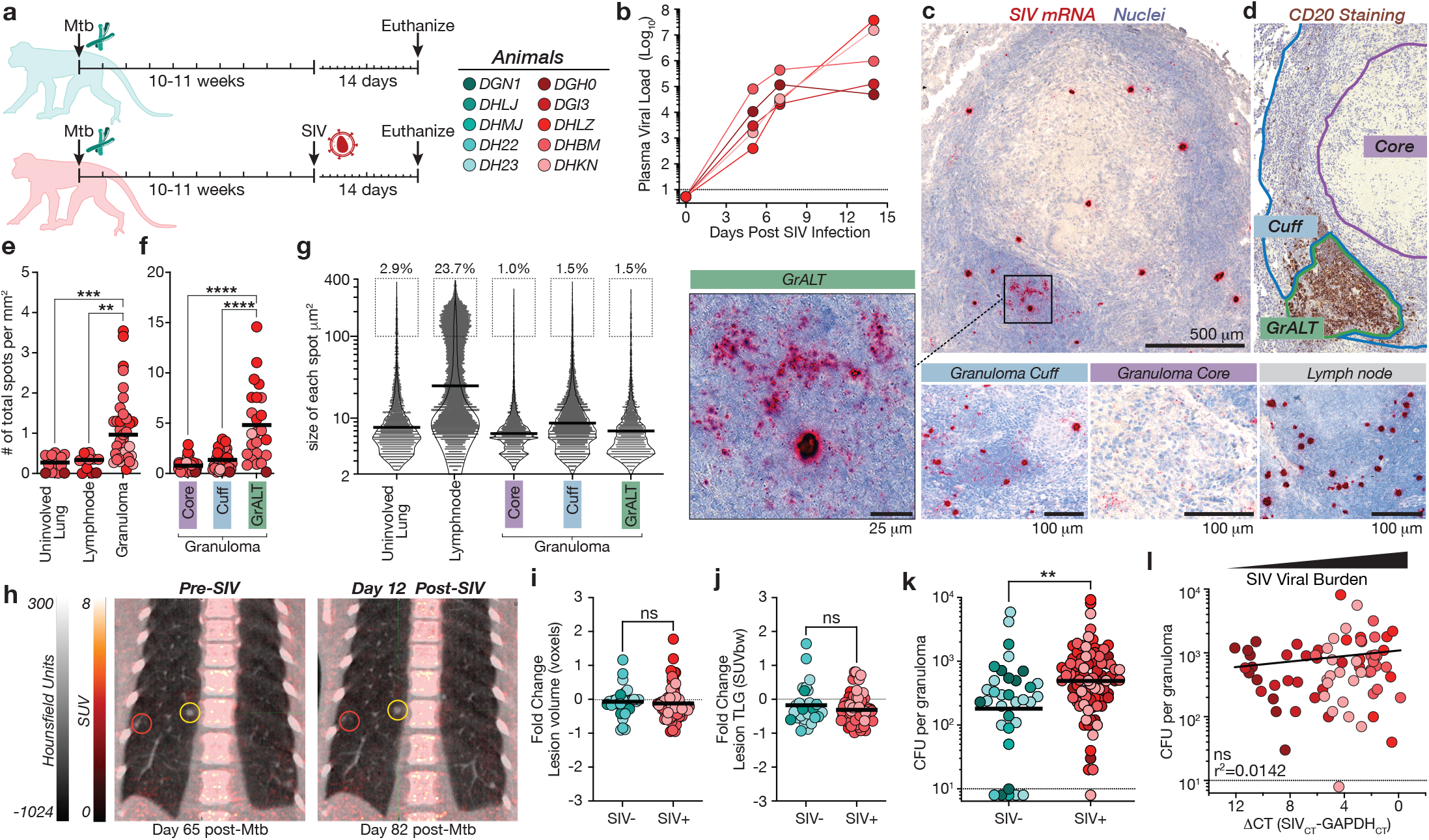
Lung Mtb granulomas are heavily infected by SIV rapidly after co-infection. (A) 10 Rhesus macaques were intrabronchially infected with Mycobacterium tuberculosis H37Rv for 10-11 weeks and 5 macaques were subsequently infected with SIVmac239 intravenously. Both groups were followed for another 14 days. (B) Acute viral infection was tracked by measuring plasma viral loads after co-infection. (C) Example image of RNAscope staining for SIV viral RNA in a co-infected granuloma. (vRNA shown in red and nuclei counterstained light blue) (D) Quantification of individual spots was performed and compared to different microanatomical locations in the granuloma as indicated and verified by immunohistochemical staining for B cell clusters. (E) Quantification of the number of SIV spots analyzed in uninvolved lung tissue and lymph nodes compared to granulomas. (F) Different regions of the granuloma were further characterized demonstrating increased viral burden in GrALT. (G) Distribution violin plots of the size in μm^2^ of individual spots with the frequency of >100 μm spots indicated above the boxed area. (H) ^18^FDG PET-CT scans were performed immediately before SIV infection and at day 12 post co-infection which allowed tracking individual lesions. (I) The change in volume of each lesion as measured in voxels (0.5 mm^3^) with with abnormal houndsfeld unit density (−400 to 200 HU) on CT was insiginifcant from pre- to post-SIV infection. (J) ^18^FDG uptake, expressed as the total lesion glycolytic (TLG) activity in standardized uptake value/body weight (SUVbw)/mL, of each lesion was compared pre- to post-SIV infection and also found to be insignificant. (K) Individual granulomas were isolated at the time of necropsy for determination of bacterial load. (L) RNA isolated from each granuloma was assayed for viral quantification and compared to bacterial burden. Statistical analysis was calculated using (E-F) one-way ANOVA, (I-J) student’s unpaired T-tests, (K) Mann-Whitney U-test, or (L) nonlinear regression analysis. ns = non-significant, ** p<0.01, *** p<0.001 **** p<0.0001

To determine if SIV co-infection led to changes in TB lesions, macaques were imaged using ^18^FDG PET-CT scanning prior to and 12 days after SIV co-infection (Fig. 1h). There were no changes in the number of lesions that could be identified by PET-CT imaging after SIV infection, however at necropsy, more granulomas were isolated from co-infected macaques (Fig. S1c). We found no impact of SIV infection on granuloma volumes as measured by volume of abnormal voxels or density range (Fig. 1i) or glycolytic activity as measured by ^18^F-FDG uptake (Fig. 1j). Although statistically significant, the difference in the size of granulomas was not notable, and while there was a trend for increased cellularity of the SIV granulomas in co-infected animals, the difference did not reach statistical significance (Fig. S1d-e). Mycobacterial loads in individual granulomas were statistically significantly increased in the co-infected animals, but the difference was less than 2 fold (~625 CFU in SIV-uninfected granulomas vs. ~955 CFU in co-infected granulomas) (Fig. 1k). Lastly, there was no significant correlation between SIV viral and mycobacterial loads in granulomas, indicating that the granulomas with lower viral loads were not different due to their mycobacterial burdens (Fig. 1k). Thus, as expected we did not observe tuberculosis disease reactivation in the first two weeks after SIV infection, however it is possible that bacterial loads were just beginning to increase and new granulomas to form.

Next, we examined the effect of acute SIV infection on the pattern of soluble markers of inflammation in granulomas. The markers that correlated with bacterial loads were different in mono-vs co-infected granulomas (Fig. 2a). Concentrations of CCL2, CCL4, IL-8, and IL-1RA were positively correlated with mycobacterial burden in the granulomas of both groups of animals. In contrast, IL-6 and IL-1 β levels only correlated with CFU in mono-infected granulomas, and CCL3, TNF, CXCL12 and GM-CSF correlated with CFU only in the SIV co-infected granulomas (Fig 2a). SIV loads in granulomas were most strongly correlated with CXCL11 (an IFN-inducible CXCR3 ligand) and to a lesser extent with CCL3 and CCL4 (both CCR5 ligands) as well as TNF, IL-18, and IL-23 (Fig 2a). SIV-infected granulomas contained higher levels of CCL3 and IL-8, but the differences were less than ~2 fold, and the granulomas from SIV infected and co-infected animals contained very similar levels of the soluble mediators measured here (Fig. 2b). Thus, although SIV infection did not dramatically impact the overall levels of soluble mediators in the first 2 weeks after infection, it altered the relationship between mycobacterial loads and inflammatory mediators in granulomas.

**Figure 2.**
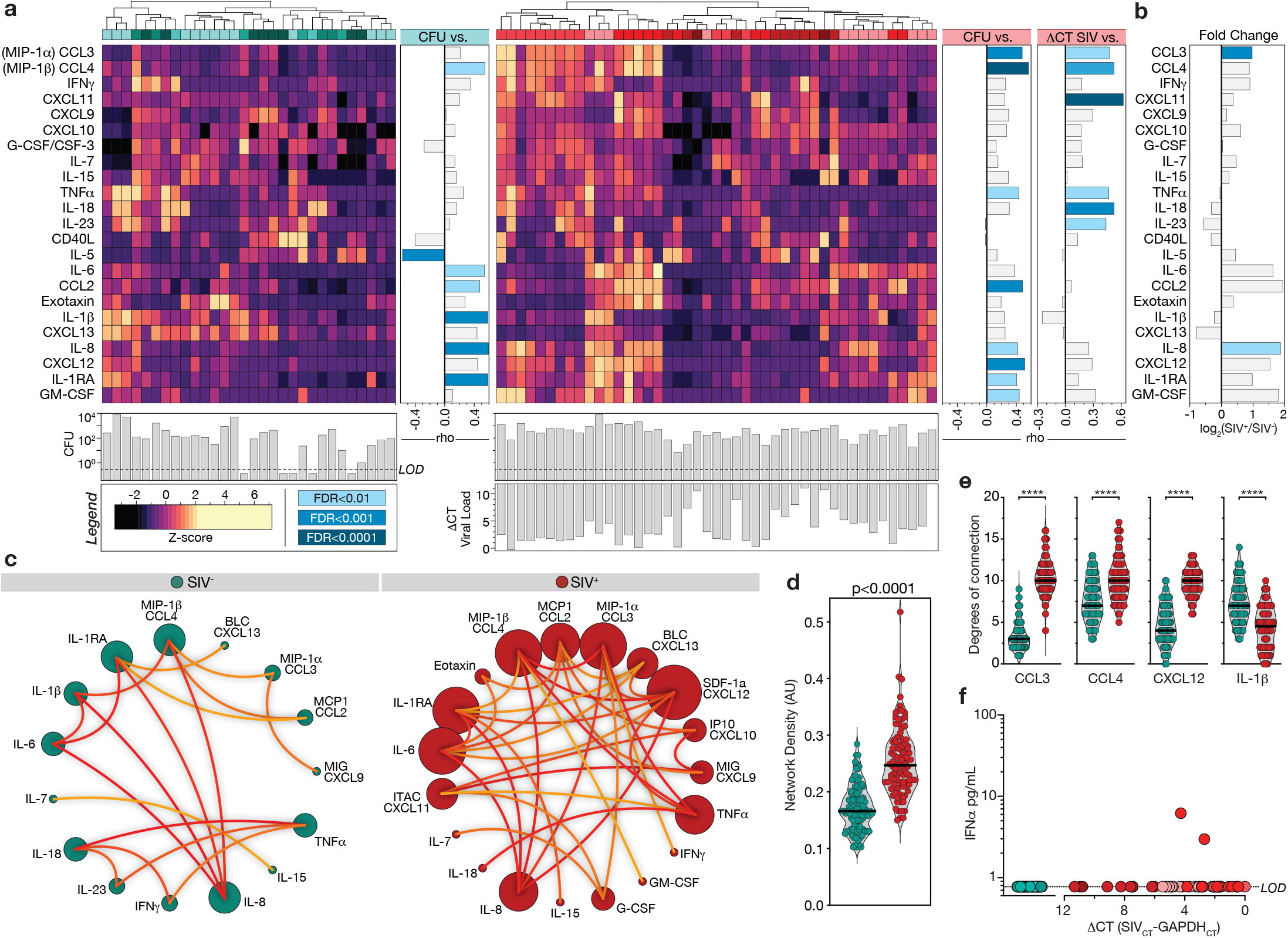
Early changes in soluble mediators in Mtb granulomas after SIV co-infection. (A) Granuloma homogenates were assayed for soluble mediators of inflammation, values normalized to protein levels and by Z-score and compared to bacterial and viral burdens. Heatmap visualization of 23 markers within mono-infected (left) and co-infected (right) granulomas with indicated bacterial burden for each granuloma shown below the heatmap, and spearman correlations with pathogen burdens to the right. Correlations were calculated intra-group and only values with false discovery rate below 0.01 highlighted as significant. (B) The log2 fold change of soluble markers between mono- and co-infected groups were calculated and compared for significance. (C) Correlation network analysis using Spearman’s rank test were performed and visualized using circus plots where the size of the circle indicates the number of significant correlations. (D) The overall distribution of network density degree and (E) individual degrees of connectivity of CCL3, CCL4, CXCL12, and IL-1β in 100 bootstrap replicates. (F) The levels of IFNa in granuloma homogenates of all samples shown in correlation to viral burden. Statistical analysis was performed using (A) Spearman’s rank test for correlations, (B, D-E) student’s T test with false discovery ratio correction, or (E) Wilcoxon signed-rank test.

To further characterize the impact of early SIV infection on the inflammatory milieu of granulomas, we next used network density analysis of Spearman correlations^24,25^ to quantify the interconnectivity of these variables (Fig. 2c). After SIV infection multiple soluble mediators, in particular several chemokines, had increased numbers of correlations with each other, and the overall network density was significantly higher in co-infected granulomas (Fig. 2d). Most notably, CXCL12 had no significant correlations with other mediators in the mono-infected granulomas but was the largest node in the co-infected granulomas. Comparing the degrees of connection from the bootstrap analysis highlighted the increased connectivity of CCL3, CCL4, and CXCL12 in co-infected granulomas while IL-1β showed increased connectivity in mono-infected granulomas (Fig. 2e). Such increased network density has previously been shown to reflect heightened inflammatory states^25–27^. Interestingly, IFNa was not detected in the granulomas from either group of animals, so we are unable to implicate virus-induced type IFN response in these changes in the granuloma inflammatory milieu after SIV infection (Fig 2f). When taken together with the overall lack of major increases in the total concentrations of most mediators, and the very small increase in bacterial loads, we speculate these data indicate that at 14 days after viral infection, SIV may be just beginning to drive increased mycobacterial loads and perturb inflammatory responses.

We next examined T cell composition after co-infection. The frequency of CD4 T cells was not significantly different across peripheral blood mononuclear cells (PBMCs), pulmonary lymph nodes (LN), splenocytes, and bronchoalveolar lavage (BAL). In contrast, CD4 T cells were greatly reduced in SIV co-infected granulomas on day 14 after SIV infection (Fig. 3a, gating strategy Fig S3). Accordingly, there was no difference in the frequency of CD8 T cells in the BAL, LN or spleen, but an increase was observed in granulomas from co-infected animals (Fig. S3d). The difference in the CD4:CD8 T cell ratio was different between *Mtb mono*-infected and *Mtb*/SIV *co*-infected granulomas (*p<0.0001*), and the decline in the CD4:CD8 cell ratio was correlated with SIV viral burden (Fig. 3b). Absolute numbers of CD4 T cells were significantly reduced in co-infected compared to mono-infected granulomas, while CD8 T cell numbers were retained (Fig. 3c). 60.5% of co-infected granulomas contained less than 20 CD4 T cells and were therefore excluded from further phenotypic analysis. MTB300 peptide-specific CD4 T cells were maintained in the pulmonary LN and BAL after SIV co-infection but were strikingly reduced in granulomas, where the majority of granulomas no longer contained any Mtb-specific CD4 T cells (Fig. 3d-e, S3e). The Mtb peptide pool used in this study is not optimized for restimulation of CD8 T cells, however, Ag-specific CD8 T cells could still be detected and were not different in any tissue sites of the mono and co-infected animals (Fig. S3f-g). Taken together these results demonstrate that CD4 T cells, and particularly Mtb-specific CD4 T cells, are depleted from the granulomas prior to detectable depletion in the peripheral tissues or even BAL, and the decline in CD4 T cells is correlated with the viral burden of individual granulomas.

**Figure 3.**
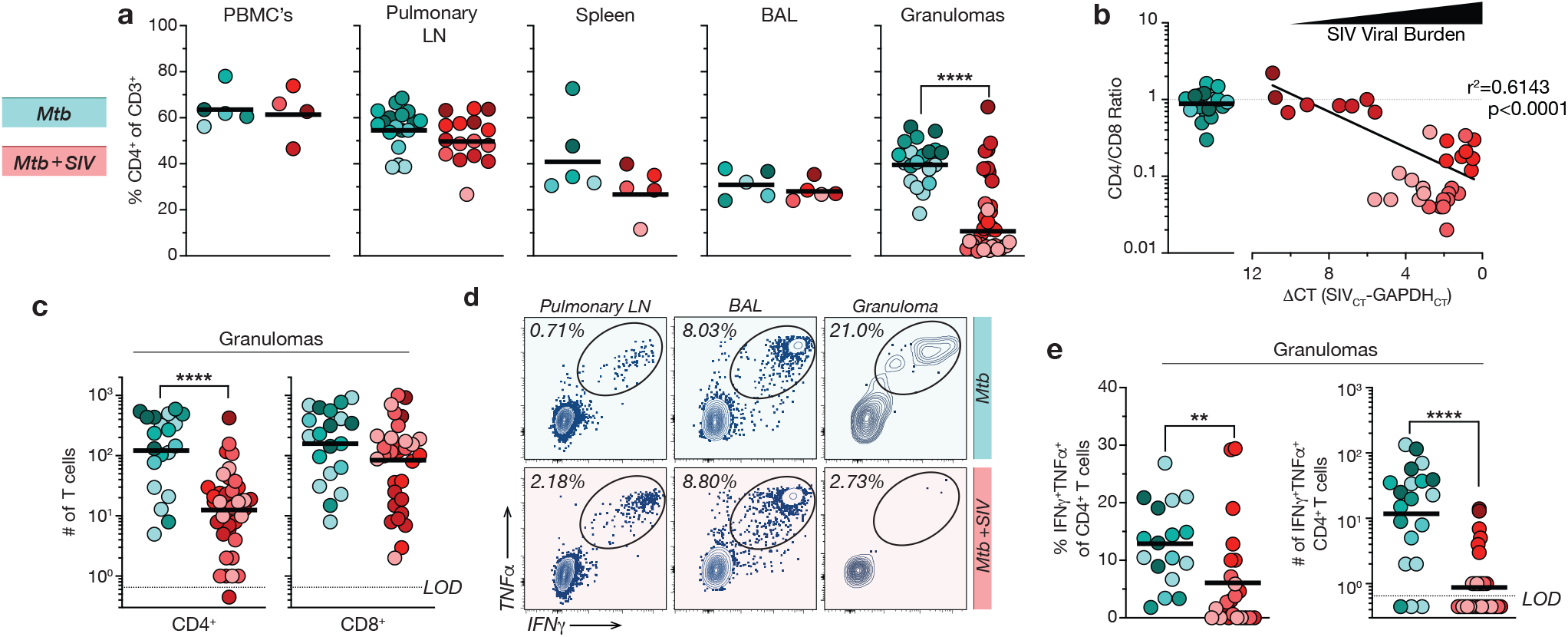
Mtb-specific CD4 T cells are massively depleted in granulomas two weeks after SIV co-infection. (A) Flow cytometric analysis of PBMC’s, pulmonary lymph nodes (LN), splenocytes, bronchoalveolar lavage (BAL), and individual granulomas showing the percentage of CD3^+^ T cells that are CD4^+^. (B) Comparison of the CD4:CD8 ratio in individual granulomas to the viral burden. (C) Quantification of the number of total CD4^+^ and CD8^+^ T cells in granulomas. LOD = Limit of detection. (D) Example flow cytometry plots of Mtb antigen stimulation of CD4^+^ T cells in pulmonary LN, BAL, and granulomas from mono- and co-infected (top and bottom row, respectively). (E) Frequency and absolute number of antigen specific CD4^+^ T cells in granulomas after restimulation with a Mtb peptide pool. Statistical analysis was calculated using (A) students unpaired T-tests, (B) nonlinear regression analysis, or (C, E) Mann-Whitney U-test. ** p<0.01, **** p<0.0001

CD4 T cells are variably susceptible to SIV-mediated depletion based on activation and differentiation state and expression viral entry co-receptors (e.g. CCR5)^14,28^. To examine the differential susceptibility of granuloma CD4 T cell subsets to SIV co-infection, we analyzed T cells for the expression of chemokine receptors CCR5, CCR6, CXCR3, CXCR5, and the transcription factor eomesodermin (eomes) (Fig. S4a). We focused on the bulk population of non-naïve (CD95^+^) CD4 T cells, due to the near complete absence of Mtb-specific CD4 T cells in most of the co-infected granulomas. There was a significant reduction in CCR5, CCR6, and CXCR3-expressing cells, no difference in eomes^+^ cells, and a slight increase in CXCR5^+^ CD4 T cells in co-infected compared to mono-infected granulomas (Fig. 4a-b). In contrast, among CD8 T cells there was an increase in CCR5^+^ and eomes^+^ cell in co-infected granulomas (Fig. 4c). Consistent with the granulomas, similar differences were also seen for both activated CD4 and CD8 T cells in the pulmonary LN (Fig. S4b-c). Boolean gating of these markers allowed us to discriminate between several relevant T cell subsets: CXCR3^+^CCR6^-^ Th1 cells, CCR6^+^CXCR3^+^ T_H_1* cells, CCR6^+^CXCR3^-^ Th17-like and CXCR5^+^ Tfh-like cells (Fig. S4a). We were able to further subdivide each of these subsets based on expression of the viral entry co-receptor CCR5, as well as eomes. This accounted for >90% of all memory T cells in the granulomas (Fig. S4d-e). Among both Th1* and Th1 cells, we found that the CCR5^+^eomes^-^ subset of these effector lineages was preferentially depleted (Fig. 4d). In contrast, both CCR5^+^eomes^+^ Th1 and Th1* cells did not display evidence of preferential depletion. In CD8 T cells, there was a significant increase in the frequency of CCR5^+^CXCR3^+^eomes^+^ T cells and commensurate reduction in CCR5^+^CXCR3^+^eomes^-^ and CCR5^-^CXCR3^+^eomes^-^ T cells from co-infected granulomas (Fig. 4e). Interestingly, the increase in CCR5^+^ CD8 T cells is consistent with the elevated levels of CCL3 we observed in granulomas. Thus, while the overall population of CD4 T cells in granulomas is greatly reduced, there is evidence of preferential depletion of the subsets of Th1 and Th1* CD4 T cells that express CCR5 but lack eomes. Future experiments are needed to explore the role of eomes in determining the susceptibility of CCR5^+^ CD4 T cells to SIV-mediated depletion.

**Figure 4.**
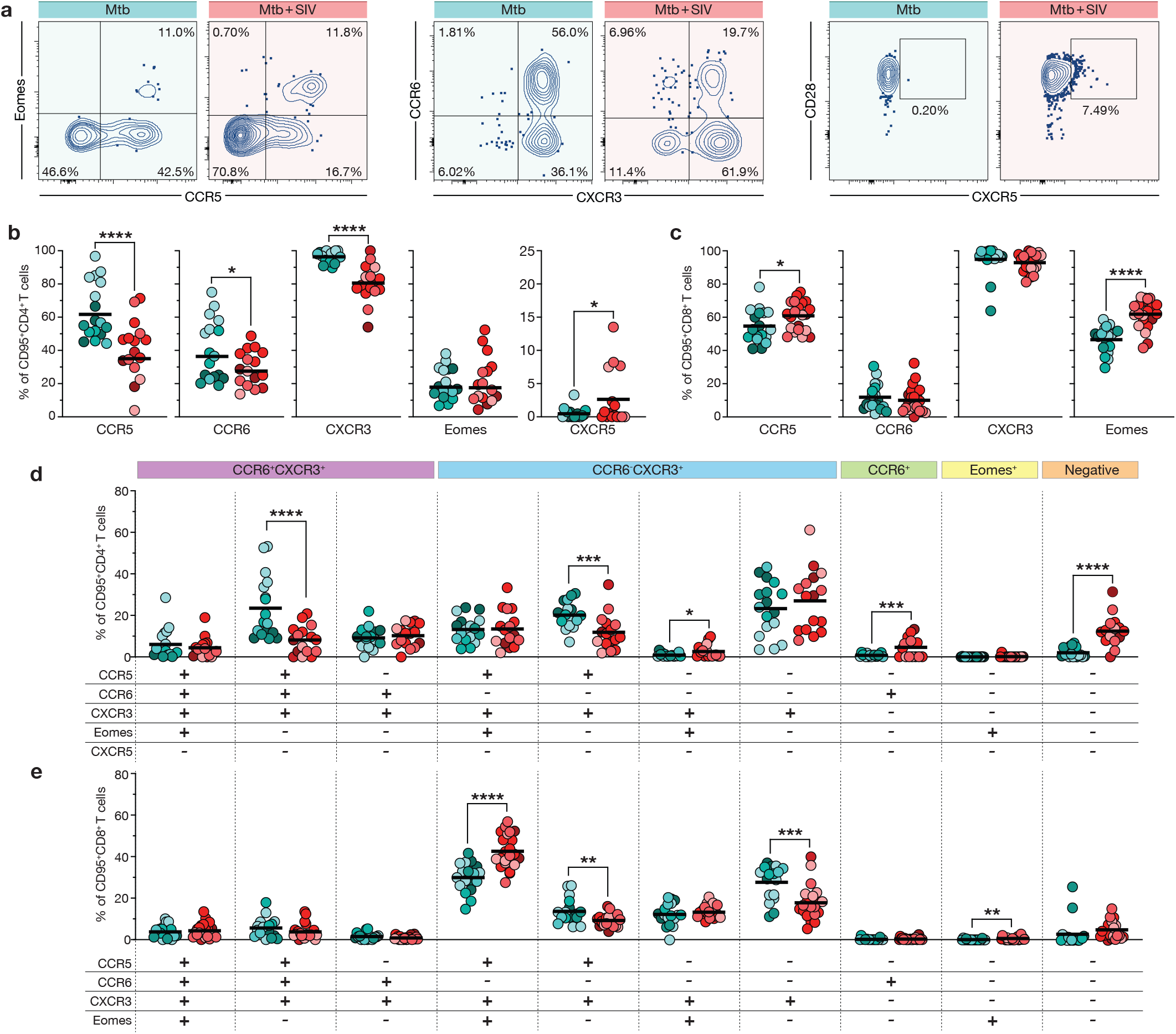
CCR5-expressing Th1 and Th1* cells in granulomas are preferentially depleted by SIV co-infection. (A) Example flow cytometry plots of activated CD4 T cells in granulomas expressing chemokine receptors CCR5, CCR6, CXCR3, CXCR5, and the transcription factor eomesodermin (eomes). (B-C) Expression of markers in non-naïve CD4 T cells and CD8 T cells. (D-E) Boolean analysis of the same markers where the ten populations of cells shown comprise >90% of total population of CD4 and CD8 T cells. Statistical analysis was calculated using Student’s unpaired T-tests. * p<0.05, ** p<0.01, *** p<0.001, **** p<0.0001

To determine the early effects of SIV infection on granuloma architecture and intralesional T cell trafficking, we performed live-imaging of granuloma thick-section explants^24^. Live sections of granulomas were stained with antibodies to CD4, CD20, and CD11b and imaged at 37°C for 1-2 hours (Fig 5a, Supplemental movie). Quality control was performed to remove tracks with less than 5 spots or whose movement was restricted by edges in the X, Y, and Z planes (Fig. S5). The X- and Y-start position of the tracks analyzed from the granuloma shown in (Fig. 5a) can be seen for CD4^+^ tracks and CD20^+^ tracks (Fig. 5b). As expected, the number of tracks normalized to the volume of tissue imaged in each granuloma was significantly lower for CD4^+^ tracks in co-infected compared to mono-infected granulomas, but there was no difference in CD20^+^ tracks between the groups (Fig. 5c). Accordingly, the ratio between CD4^+^ and CD20^+^ tracks was statistically significantly reduced in granulomas imaged from co-infected macaques (Fig. S6a). We next quantified the localization of tracks within the macrophage-rich core, lymphocyte-rich cuff, or B cell-rich GrALT regions of granulomas (Fig. 5b). In SIV co-infected granulomas, the frequency of CD4^+^ tracks localized to GrALT was increased due to a reduction in the absolute number of tracks found in the core and cuff (Fig. 5d-e). In contrast, the distribution of CD20^+^ tracks to different regions of the granuloma was not different between mono- and co-infected granulomas (Fig. S6b-c). These results demonstrate CD4 T cells in the core and cuff (i.e., CD4 T cells best positioned to interact with Mtb-infected macrophages), are preferentially depleted early after SIV infection.

**Figure 5.**
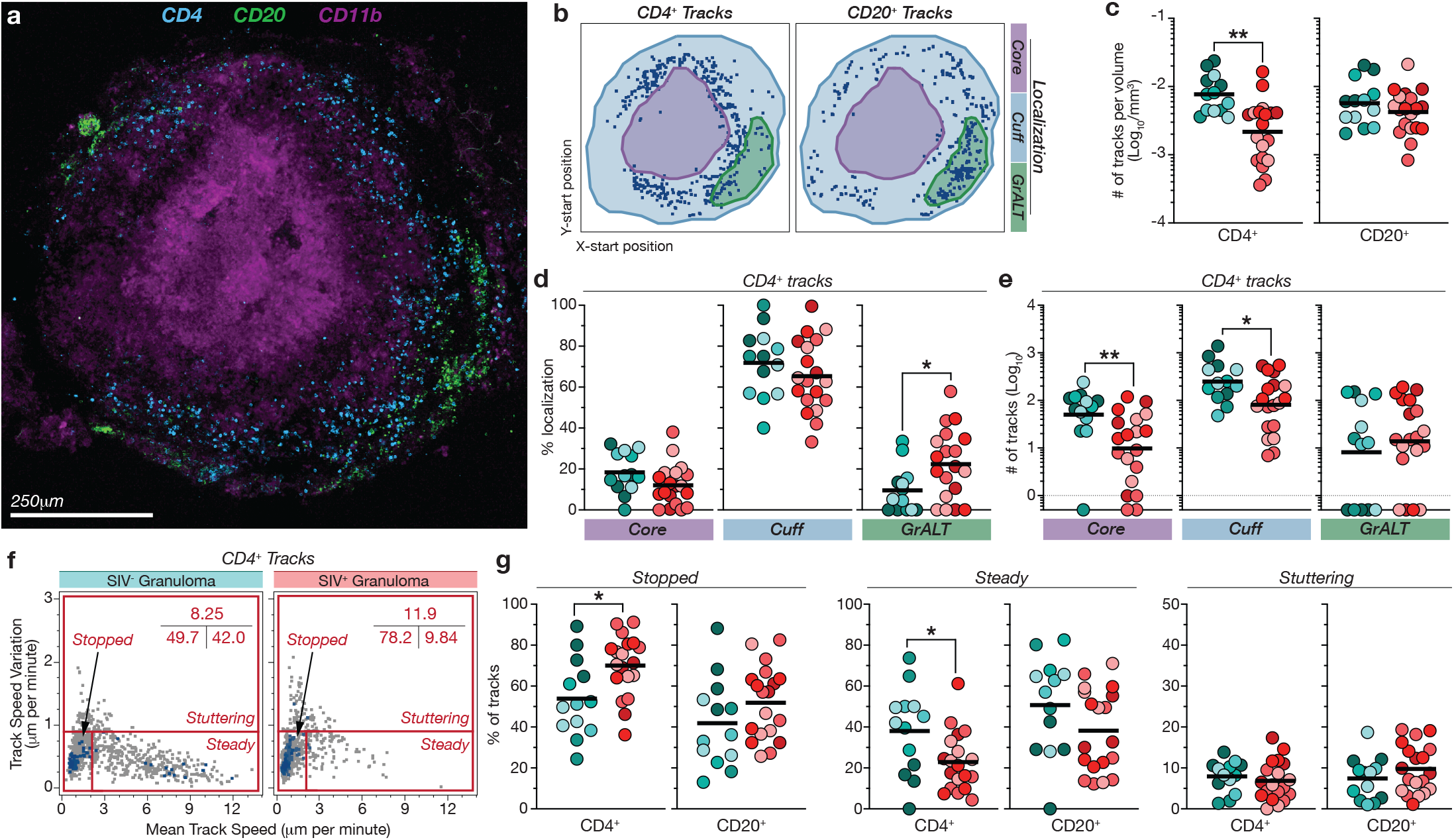
SIV co-infection alters the spatial distribution and intralesional trafficking of CD4 T cells in Mtb granulomas. (A-B) Example still image of a granuloma used for 4D imaging (CD4 is cyan, CD20 is green, and CD11b is purple) where distilled cellular tracks from CD4^+^ and CD20^+^ cells shown in the X- and Y-start position. (C) Absolute number of tracks normalized to the volume of tissue imaged. (D-E) Localization of CD4^+^ tracks within the macrophage rich core, the lymphocyte dense cuff, or the granuloma-associated lymphoid tissue that forms within the cuff as the frequency of or absolute number of total CD4^+^ tracks in the granuloma. (F) Example plots of the movement of CD4^+^ tracks from a single granuloma with frequency denoted in the top right corner. (G) Quantification of the movement of CD4^+^ tracks and CD20^+^ tracks as denoted as stopped, steadily moving, or stuttering. Statistical analysis was calculated using students unpaired T-tests. * p<0.05, ** p<0.01

We next compared CD4 T cell and B cell movement in mono- and co-infected granulomas. Individual tracks were categorized based on their mean speed and the variation in speed into 3 major patterns of cellular motility: stopped cells, stuttering cells, or steadily moving cells (Fig. 5f). CD4^+^ tracks displayed reduced motility evidenced by a significant increase in stopped tracks and concordant decrease in steadily moving tracks (Fig. 5g). This difference was not seen in CD20^+^ tracks where there was no difference stopped, steady, or stuttering tracks in mono- vs co-infected granulomas (Fig. 5g). This difference was even more significant when pairing the analysis of CD4^+^ and CD20^+^ tracks from the same granuloma indicating the changes in motility are not granuloma specific but CD4 T cell specific (Fig. S6d). Taken together these results demonstrate that SIV co-infection significantly decreased CD4 T cell movement in granulomas.

## DISCUSSION

CD4 T cell depletion is considered the primary mechanism by which HIV infection leads to the inability to control growth of Mtb, but little is understood about the impact of HIV infection on CD4 T cells at the sites of mycobacterial infection in the tissues. Here we used a non-human primate model of TB and SIV co-infection to study the early events of SIV infection on CD4 T cells in Mtb granulomas. We find that CD4 T cells in granulomas are highly sensitive to SIV-mediated depletion compared to other tissues. It is not clear why granuloma CD4 T cells are so susceptible to SIV-mediated depletion. However, it is reminiscent of the massive loss of CD4 T cells in the gastrointestinal tract of PLWH and of non-human primate species that develop AIDS after SIV infection^28–31^. Moreover, lung parenchymal CD4 T cells are more susceptible than BAL CD4 T cells to HIV mediated depletion in a humanized mouse model of HIV infection^32^. The selective loss of granuloma CD4 T cells may be due to enhanced permissiveness of these T cells to SIV infection as compared to CD4 T cells in other locations (e.g. CCR5-expressing T cells are enriched in granulomas)^33^. It is also possible that the microenvironment the granuloma itself may facilitate depletion of CD4 T cells. For example, activated macrophages efficiently take up infected CD4 T cells^34^, and granulomas are densely packed with macrophages. Importantly, these two possibilities are not mutually exclusive.

In some granulomas, there was a near total loss of CD4 T cells within 14 days, indicating that most or maybe all T cells in granulomas are susceptible to depletion during SIV co-infection. Nonetheless, our data also indicate that some CD4 T cells in granulomas are even more susceptible than others. Previous reports have shown that in persons latently infected with Mtb HIV co-infection preferentially infects and depletes Mtb-specific CD4 T cells^14,35–38^. Perhaps most importantly, our data show that Mtb-specific CD4 T cells were greatly reduced in frequency in SIV co-infected granulomas long before depletion manifested in blood. CCR5-expressing Th1 and Th1* cells were also reduced in frequency in SIV co-infected granulomas. This is expected as CCR5 is a co-receptor for viral entry. Interestingly, we found that CCR5^+^ cells that also expressed the transcription factor eomes were not reduced in frequency in co-infected lesions. The mechanisms underlying this apparent differential sensitivity of CCR5^+^eomes^+^ CD4 T cells is not clear. Expression of CCR5 ligands in CD4 T cells has been shown previously to be associated decreased viral replication in viremic patients and elite controllers^39,40^, and eomes^+^ CD4 T cells have been shown to produce high levels of CCL3/MIP-1a^41^, which may partly explain the increase of soluble CCL3 in granulomas. Therefore, the preservation of CCR5^+^eomes^+^ CD4 T cells in granulomas may reflect protection by CCR5-ligand production.

We observed a regionalized depletion where CD4 T cells were lost to a lesser degree in GrALT as compared to the core and cuff of the granuloma. This was not due to a lack of viral replication, as the highest amounts of virus in the granuloma as detected by RNAscope was in GrALT structures. Germinal centers have been proposed to be an ‘immunologically privileged’ site for viral persistence, perhaps due to the inability of cytotoxic T cells to interact with infected T_FH_ CD4 T cells, so it is possible that less effective killing by SIV-specific cytotoxic T cells may be responsible for the decreased depletion of GrALT-resident CD4 T cells^42,43^.

We also found that the CD4 T cells remaining in SIV co-infected granulomas demonstrated reduced motility, raising the possibility that SIV-mediated suppression of CD4 T cell movement is another mechanism of immunosuppression. HIV infection has been shown to result in reduced motility of CD4 T cells in response to chemokines in vitro^44,45^. We should additionally point out, however, that CD4 T cells in granulomas may also stop moving as a result of interaction with peptide-presenting cells^46^, so the importance of this observation to the loss of bacterial control is not clear. Nonetheless, CD4 T cell interaction with infected macrophages is necessary for control of intracellular mycobacterial growth^47,48^, and it is possible that reduced trafficking of the few remaining CD4 T cells in the granulomas further impairs microbial immunity. We do not know if this defect in T cell motility occurs in other tissues, as we have only imaged lung granulomas for this work. Future studies are needed to address the broader possibility that lentiviral infection results in immunosuppression not only by killing T cells but also by limiting the mobility the ones that remain.

Collectively, these data show how SIV infection is even more detrimental for Mtb-specific immunity than previous appreciated. We should point out that we do not exclude other mechanisms of viral infection-enhanced susceptibility to Mtb infection. Indeed, cytomegalovirus infection in humans has been associated with enhanced risk of TB^49,50^ and influenza infection in mice has been shown to enhance Mtb infection^51^. SIV preferentially kills Mtb-specific CD4 T cells in granulomas, CD4 T cells phenotypically most correlated with immune control, and CD4 T cells in the subregions of granulomas that are best positioned to interact with Mtb-infected macrophages. To make things worse, it impairs the trafficking of the CD4 T cells that remain. Most patients are just beginning to notice signs and symptoms of HIV infection 14 days after primary infection, yet here we demonstrate that CD4 T cells within the granuloma have already been depleted. These data provide a potential explanation of the increased risk of TB reactivation prior to the loss of peripheral CD4 T cells in HIV infection. Understanding the granuloma-specific mechanisms CD4 T cell susceptibility to SIV-mediated depletion may lead to intervention strategies to preserve anti-tuberculosis immunity in co-infected individuals.

## METHODS

### Study Design and Infections

Rhesus macaques originally from the Morgan Island NIH breeding colony were housed in biocontainment according to the Animal Welfare Act and the Guide for the Care and Use of Laboratory Animals within a AAALAC international-accredited animal biosafety level-3 vivarium. Daily enrichment was provided for the macaques. All procedures were performed using anesthetics according to the approved NIAID DIR Animal Care and Use Committee study proposal LPD-25E. Additionally, animals were scanned at regular intervals with a LFER 150 PT/CT scanner (Mediso, Inc., Hungary) using ^18^FDG (0.5 mCi/kg) and tubercular lung disease, as well as, individual lesion characteriztics were analyzed on serial scans as described previously^24,52^. The object of this study was to understand the effects of acute SIV on Mtb-specific immune responses. To achieve this, 10 macaques were intrabronchially infected with ~56 CFU of *Mycobacterium tuberculosis* (*Mtb*) strain H37Rv-mCardinal (kindly provided by Dr. Clifton Barry III, chief of the Tuberculosis Research Section, Laboratory of Clinical Immunology and Microbiology, NIAID) bilaterally using 2-mL of saline each into the right and left lower lobes (14.0 ± 3.77 CFU/mL). After *Mtb* infection, animals were montored for clinical signs of tuberculosis such as weight loss and pyrexia for 10-11 weeks. Thereafter animals were randomly assigned to one of two groups where the *mono*-infection group remained SIV-naïve and the co-infection group was intravenously injected with 3,000 TCID_50_ of SIVmac239. For tissue collection exactly 14 days post SIV infectioin, a co-infection animal along with a time-matched *Mtb* mono-infected animal was euthanized according to American Veterinary Medical Association guidelines.

### Tissue Processing for Cells, Bacteria, and Virus

Blood was collected using EDTA-tubes and cells isolated using Ficoll-Paque Density Centrifugation (GE Life Sciences, Canada). Bronchoalveolar lavage (BAL) was collected using a modified feeding tube inserted through the trachea into the lower pulmonary lobes and lavaged using ~60mL of sterile saline. Fluid was then filtered using a 100-micron filter and centrifuged for the collection of cells. Tissues were taken for analysis at the time of necropsy. For single cell suspensions, spleen and lymphnodes were homogenized using gentleMACSs dissociators (Miltenyi Biotec, Germany) and filtered through a 100-micron filter while isolated granulomas were mashed through a 100-micron filter using a syringe plunger. Homogenates from granulomas and tissues were serially diluted and plated on 7H11 agar plates incubated at 37°C for 3-4 weeks before quantification. Granuloma homogenate was taken for RNA isolation using RNeasy Plus Kits and stored at −80°C (Qiagen, USA). Quantification of viral burden was done using TaqPath 1-step RT-qPCR Master Mix (Thermo Fisher Scientific, USA) and primers/probes sets targeting SIVmac239 (Eurofins Scientific, USA) or GAPDH (Thermo Fisher Scientific, USA). Reactions were completed in triplicates for 40 cycles using FAM dyes. Plasma was filter-sterilized and analyzed for plasma viral load by RT-qPCR by Quantitative Molecular Diagnostics Core at the Frederick National Laboratory.

### Cell stimulation and Flow cytometry

Cells were stimulated with 2 μg/mL of MTB300 epitope peptide pool (A&A Labs, USA) in the presence of brefeldin A and monensin (Thermo Fisher Scientific, USA) for 6 hours at 37°C in Complete media (RPMI-1640 with 10% fetal calf serum (FCS), 1% Sodium pyruvate, 1% penicillin and streptomycin, 25 mM Hepes, and 2 mM l-glutamate). Fresh cells were incubated in blocking buffer containing 1% FCS, 1 μg/mL Human FC block (BD Biosciences, USA) for 1-6 hours at 4°C. Cells were then stained using fixable live/dead stain for 20 minutes at room temperature. Surface antibody staining was done in PBS containing 1% FCS and 1X Brilliant Stain Buffer (BD Biosciences, USA) for 20 minutes at room temperature. After fixation and permeabilization using Foxp3/Transcription Factor Staining Kit (eBiosciences, USA), intracellular antibody staining was done in PBS containing 1% FCS, 1X Brilliant Stain Buffer, and 1X Permeabilization reagent for 30 minutes at 4°C. Samples were acquired using a FACSymphony cytometer (BD Biosciences, USA) and data analyzed using FlowJo 10 (BD Biosciences, USA). Gating strategies are shown in figures S3-4.

### Multiplex cytokine analysis

Granuloma homogenates were filter-sterilized and processed for soluble protein marker concentrations using the Invitrogen Cytokine and Chemokine 30-Plex ProcartaPlex Kit (Thermo Fisher Scientific, USA) according to manufacturer protocol. Samples were acquired using a MAGPIX with xPONENT software (Luminex Corporation, USA). Soluble marker protein levels were normalized to total protein levels measured by Quant-iT protein Assay Kit (Thermo Fisher Scientific, USA).

### 4D Confocal Imaging of Granulomas

Isolated granulomas were kept on ice until embedding in PBS with 2% agarose. The tissue was cut into 300-micron thick sections using a Leica VT1000 S Vibrating Blade Microtome (Leica Microsystems, USA) housed in Class II, Biosafety hood within a BSL3 laboratory. Sections of tissue were incubated in 1X PBS supplemented with 10% FCS, isotype-specific blocking antibodies, and Human FC Block (BD Biosciences, USA) for 2-8 hours at 4°C. After blocking, tissues were stained with antibodies CD4, CD20, and CD11b for 2-12 hours at 4°C with intermittent shaking. Tissues were then washed and placed into chamber slides containing Complete Imaging Media (Complete media made with phenol red-free RPMI) supplemented with ProLong Live Antifade Reagent (Thermo Fisher Scientific, USA). After incubation in a 5% CO_2_ incubation chamber at 37°C for 1 hour, the chambered slide was imaged using Leica SP5 inverted confocal microscope with environmental chamber to maintain the sample humidified at 37°C (Leica Microsystems, USA). Sections were serially imaged over the course of 1-2 hours and compiled using LAS X software (Leica Microsystems, USA). Image analysis was performed using Imaris software (Bitplane, Switzerland) for quantification of individual cells using the spot function to track their movement over time. Data from tracks were exported, processed through a custom Python 3 script to select the parameters to include in the exported files, and then imported into FlowJo 10 software (BD Biosciences, USA) for data visualization and analysis as described in Fig S6. Briefly, tracks that were limited in movement by the edges of the X-, Y-, or Z-plane were excluded from analysis. Using the X- and Y-starting point of a track, cells could be localized to different areas in the granuloma. Finally, track movement was categorized into three different forms. Stopped and Steady tracks both had ≤0.90 μm/minute track speed variation but were characterized by ≤2.0 μm/minute or >2.0 μm/minute mean track speed, respectively. Stuttering tracks had any mean track speed but were characterized by track speed variation >0.90 μm/minute.

### RNAscope and Image Analysis

Formalin Fixed Paraffin embedded tissues sections were cut into 10-micron thick sections using RNAse precautions and used for RNAscope *in situ* hybridization staining for viral RNA (ACD Biotechne, USA). Slides were dewaxed using xylene and ethanol and then underwent epitope retrieval using heat induced low-pH methods according to the manufacturer. Slides were then treated with diluted protease plus for 20 min at 40°C and endogenous peroxidases blocked using hydrogen peroxide for 10 min at room temperature. Probes for SIVmac239, containing 83 separate pairs of probes spanning the proviral gag, vif, pol, tat, env, vpx, vpr, nef, and rev genes, were used for single color immunohistochemistry with hematoxylin counter-staining using the RNAscope 2.5 HD Assay-RED (ACD Biotechne, USA). Immunocytochemically stained slides were imaged using Aperio VERSA (Leica Microsystems, USA) and analyzed using a custom quPath script to identify spots with pixel size 0.5-microns, background radius 8-microns, median radius 8-microns, sigma 0.75-microns, minimum area 2.0-microns, maximum area 400-microns, with threshold of 0.25. Spots were then colocalized to granuloma regions using annotation tabs for each area of the granuloma which were identified according to the different hematoxylin counter-staining pattern and CD20 staining.

### Statistical analyses

All statistical analyses were conducted using GraphPad Prism V8 (GraphPad Software, USA) except for when specified otherwise. Data were tested for Gaussian distribution using the DÁgostino’s K-square test. For group comparisons, individual tests vary and are denoted in each figure legend but include two-tailed unpaired T-tests, Mann-Whitney *U* (when parameter compared were not normally distributed), One-way ANOVA with correction for multiple tests using Tukey Multiple Comparison test corrections, non-linear regression analysis of semi-log line, or simple linear regression analysis. Multiplex Luminex assay data were analyzed using R 4.0.2 packages *igraph, ggplot2, viridis, ComplexHeatmap, Hmisc,* with *p.adjust* function. Fold change was measured and significance test by Student’s T test with False Discovery Ratio correction. Correlation analyses were performed using the Spearman’s rank test with 100 bootstrap replicates. Bootstrap threshold value was defined in 80 replicates. Significant values were corrected using False Discovery Ratio. Any place p-value are reported in asterisks * p<0.05, ** p<0.01, *** p<0.001, and **** p<0.0001, ns=non-signficant.

---

**Supplemental Figure 1. Clinical parameters in mono- and co-infected macaques.** (A-B) Weight change as percent deviation from starting weight and rectal temperature over the course of the study. (C) Quantification of the number of visible lesions by PET/CT imaging at pre- and post-SIV viral infection along with the number of lesions isolated at the time of necropsy. (D) The diameter of individual granulomas measured at necropsy. (E) Number of cells isolated per granuloma after processing for single cell suspensions. Statistical analysis was calculated using (C) paired Student’s T test or (D-E) Mann-Whitney U-test. ns = not significant

**Supplemental Figure 2. RNAscope imaging quantification of SIV RNA**. (A-B) RNAscope image of a SIV-infected granuloma with SIV viral RNA shown in red with a light blue nuclei counterstain with higher-magnification image inset. (C-D) Quantification of viral spots using QuPath Image analysis, shown as yellow spots with the same high-magnification image shown above. (E) Example gating pattern for the localization of spots throughout the granuloma. (F) Image of SIV-naïve granuloma to demonstrate the high specificity of in situ hybridization.

**Supplemental Figure 3. T cell gating strategy.** Shown is an example flow cytometric gating strategy for a single cell suspension from a pulmonary LN. (A) Parent gates for lymphocytes, live cells, and singlets with progression to CD3^+^ T cells and frequency of CD4^+^ to CD8^+^ T cells. Detailed analysis of T cells starting from parent gates and progressing to (B) CD3^+^CD4^+^CD8^-^ CD95^+^ T cells or (C) CD3^+^CD4^-^ CD8^+^CD95^+^ T cells. (D) Frequency of CD3^+^ T cells that are CD8^+^ in the blood, pulmonary LN, spleen, and BAL. (E) Frequency of Mtb-specific CD4^+^ T cells in the pulmonary LN and BAL at the time of necropsy. (F) Example flow cytometry plots of CD8^+^ T cells stimulated with Mtb peptide pools (14-mer epitopes) and the (G) quantification of antigen-specific CD8^+^ T cells in granulomas, BAL, and pulmonary LN’s. Statistical analysis was calculated using Student’s unpaired T-tests. **** p<0.0001

**Supplemental Figure 4. Differential depletion of phenotypically distinct CD4 T cell subsets.** (A) Example gating strategy used for the Boolean analysis of CCR5, CCR6, CXCR3, CXCR5, and eomes in both CD4^+^ and CD8^+^ T cells. Parent gate represents gating strategy shown in Supplemental Figure 3 as labelled above initiating plots. (B-C) Expression of these markers on activated CD4 and CD8 T cells from the pulmonary LN. (D-E) Quantification of the percent of total CD4 and CD8 T cells comprised of the 10 populations shown in Fig. 4D-E. Statistical analysis was calculated using students unpaired T-tests. * p<0.05, **** p<0.0001

**Supplemental Figure 5. Analysis strategy for 4D imaging of granulomas.** (A) Example plots of the stringent parameters used to isolate CD4 and CD20 tracks. Only tracks with more than or equal to 5 individual spots were used, with cells excluded that were inhibited by the Z, X, or Y boundaries. (B) The tracks analyzed for this granuloma with comparison of a still-image. (C) High-magnification image shows spots analyzed overlayed with the actual image. (D) Example flow plots of cellular movement in two different granulomas showing the range of movement and indicated movement characteristics.

**Supplemental Figure 6. Analysis of cellular movement in granulomas.** (A) Ratio of CD4 to CD20 tracks in all granulomas analyzed by 4D imaging. (B-C) Quantification of the frequency and absolute number of CD20 tracks from the core, cuff, and GrALT. (D) Paired analysis of CD4 and CD20 tracks from the same granuloma to show the reduction in movement as specific to T cells and not the whole granuloma. Statistical analysis was calculated using (A-C) students unpaired T-tests and (D) paired T-tests. * p<0.05, *** p<0.001, **** p<0.0001

## Acknowledgements

We are very grateful to the veterinary care provided by Dr. Rashida Moore and the NIAID, Comparative Medicine Branch Animal Biosafety Level 3 facility staff. We also thank Drs. Alan Sher and Eduardo P. Amaral for discussion of data and feedback on the manuscript. We would like to thank the members of the NIAID/DIR Tuberculosis Imaging Program for their assistance: Ayan Abdi, Joel D. Fleegle, Felipe Gomez, Michaela K. Piazza, Katelyn M. Repoli, Becky Y. Sloan, Ashley L. Butler, April M. Walker, Danielle M. Weiner, Michael J. Woodcock, and Alexandra Vatthauer.

The contents of this publication do not necessarily reflect the views or policies of the Department of Health and Human Services, nor does mention of trade names, commercial products, or organizations imply endorsement by the United States government. The authors declare no conflict of interest.

## REFERENCES

1. Uncategorized References

1. Geldmacher, C., Zumla, A. & Hoelscher, M. Interaction between HIV and *Mycobacterium tuberculosis:* HIV-1-induced CD4 T-cell depletion and the development of active tuberculosis. Curr Opin HIV AIDS 7, 268–275 (2012).

2. Okoye, A.A. & Picker, L.J. CD4(+) T-cell depletion in HIV infection: mechanisms of immunological failure. Immunol Rev 254, 54–64 (2013).

3. Glynn, J.R., et al. Effects of duration of HIV infection and secondary tuberculosis transmission on tuberculosis incidence in the South African gold mines. AIDS 22, 1859–1867 (2008).

4. van der Sande, M.A., et al. Incidence of tuberculosis and survival after its diagnosis in patients infected with HIV-1 and HIV-2. AIDS 18, 1933–1941 (2004).

5. Esmail, H., et al. The Immune Response to *Mycobacterium tuberculosis* in HIV-1-Coinfected Persons. Annu Rev Immunol 36, 603–638 (2018).

6. Global Tuberculosis Report 2019, (Geneva: World Health Organization, 2019).

7. Mayer-Barber, K.D., et al. Host-directed therapy of tuberculosis based on interleukin-1 and type I interferon crosstalk. Nature 511, 99–103 (2014).

8. Zhang, L., Jiang, X., Pfau, D., Ling, Y. & Nathan, C.F. Type I interferon signaling mediates *Mycobacterium tuberculosis-induced* macrophage death. J Exp Med 218(2021).

9. Sandler, N.G., et al. Type I interferon responses in rhesus macaques prevent SIV infection and slow disease progression. Nature 511, 601–605 (2014).

10. DiNapoli, S.R., Hirsch, V.M. & Brenchley, J.M. Macrophages in Progressive Human Immunodeficiency Virus/Simian Immunodeficiency Virus Infections. J Virol 90, 7596–7606 (2016).

11. Kuroda, M.J., et al. High Turnover of Tissue Macrophages Contributes to Tuberculosis Reactivation in Simian Immunodeficiency Virus-Infected Rhesus Macaques. J Infect Dis 217, 1865–1874 (2018).

12. Brenchley, J.M., et al. CD4+ T cell depletion during all stages of HIV disease occurs predominantly in the gastrointestinal tract. J Exp Med 200, 749–759 (2004).

13. Picker, L.J., et al. Insufficient production and tissue delivery of CD4+ memory T cells in rapidly progressive simian immunodeficiency virus infection. J Exp Med 200, 1299–1314 (2004).

14. Geldmacher, C., et al. Preferential infection and depletion of *Mycobacterium tuberculosis*-specific CD4 T cells after HIV-1 infection. J Exp Med 207, 2869–2881 (2010).

15. Pagan, A.J. & Ramakrishnan, L. The Formation and Function of Granulomas. Annu Rev Immunol 36, 639–665 (2018).

16. Mattila, J.T., et al. Microenvironments in tuberculous granulomas are delineated by distinct populations of macrophage subsets and expression of nitric oxide synthase and arginase isoforms. J Immunol 191, 773–784 (2013).

17. Kauffman, K.D., et al. Defective positioning in granulomas but not lung-homing limits CD4 T-cell interactions with *Mycobacterium tuberculosis-infected* macrophages in rhesus macaques. Mucosal Immunol 11, 462–473 (2018).

18. Ulrichs, T., et al. Human tuberculous granulomas induce peripheral lymphoid follicle-like structures to orchestrate local host defence in the lung. J Pathol 204, 217–228 (2004).

19. Phuah, J.Y., Mattila, J.T., Lin, P.L. & Flynn, J.L. Activated B cells in the granulomas of nonhuman primates infected with *Mycobacterium tuberculosis*. Am J Pathol 181, 508–514 (2012).

20. Foreman, T.W., et al. CD4+ T-cell-independent mechanisms suppress reactivation of latent tuberculosis in a macaque model of HIV coinfection. Proc Natl Acad Sci U S A 113, E5636–5644 (2016).

21. Diedrich, C.R., et al. SIV and *Mycobacterium tuberculosis* synergy within the granuloma accelerates the reactivation pattern of latent tuberculosis. PLoS Pathog 16, e1008413 (2020).

22. Larson, E.C., et al. Pre-existing Simian Immunodeficiency Virus Infection Increases Expression of T Cell Markers Associated with Activation during Early *Mycobacterium tuberculosis* Coinfection and Impairs TNF Responses in Granulomas. J Immunol (2021).

23. Ganatra, S.R., et al. Antiretroviral therapy does not reduce tuberculosis reactivation in a tuberculosis-HIV coinfection model. J Clin Invest 130, 5171–5179 (2020).

24. Kauffman, K.D., et al. PD-1 blockade exacerbates *Mycobacterium tuberculosis* infection in rhesus macaques. Sci Immunol 6(2021).

25. Vinhaes, C.L., et al. Dissecting disease tolerance in *Plasmodium vivax* malaria using the systemic degree of inflammatory perturbation. PLoS Negl Trop Dis 15, e0009886 (2021).

26. Tiburcio, R., et al. Dynamics of T-Lymphocyte Activation Related to Paradoxical Tuberculosis-Associated Immune Reconstitution Inflammatory Syndrome in Persons With Advanced HIV. Front Immunol 12, 757843 (2021).

27. Bruyn, E.D., et al. Inflammatory profile of patients with tuberculosis with or without HIV-1 co-infection: a prospective cohort study and immunological network analysis. Lancet Microbe 2, e375–e385 (2021).

28. Okoye, A., et al. Progressive CD4+ central memory T cell decline results in CD4+ effector memory insufficiency and overt disease in chronic SIV infection. J Exp Med 204, 2171–2185 (2007).

29. Veazey, R.S., et al. Identifying the target cell in primary simian immunodeficiency virus (SIV) infection: highly activated memory CD4(+) T cells are rapidly eliminated in early SIV infection in vivo. J Virol 74, 57–64 (2000).

30. Mattapallil, J.J., et al. Massive infection and loss of memory CD4+ T cells in multiple tissues during acute SIV infection. Nature 434, 1093–1097 (2005).

31. Brenchley, J.M., et al. Differential Th17 CD4 T-cell depletion in pathogenic and nonpathogenic lentiviral infections. Blood 112, 2826–2835 (2008).

32. Corleis, B., et al. HIV-1 and SIV Infection Are Associated with Early Loss of Lung Interstitial CD4+ T Cells and Dissemination of Pulmonary Tuberculosis. Cell Rep 26, 1409–1418 e1405 (2019).

33. Paiardini, M., et al. Low levels of SIV infection in sooty mangabey central memory CD(4)(+) T cells are associated with limited CCR5 expression. Nat Med 17, 830–836 (2011).

34. Calantone, N., et al. Tissue myeloid cells in SIV-infected primates acquire viral DNA through phagocytosis of infected T cells. Immunity 41, 493–502 (2014).

35. Bunjun, R., et al. Th22 Cells Are a Major Contributor to the Mycobacterial CD4(+) T Cell Response and Are Depleted During HIV Infection. J Immunol 207, 1239–1249 (2021).

36. Strickland, N., et al. Characterization of *Mycobacterium tuberculosis-Specific* Cells Using MHC Class II Tetramers Reveals Phenotypic Differences Related to HIV Infection and Tuberculosis Disease. J Immunol (2017).

37. Geldmacher, C., et al. Early depletion of *Mycobacterium tuberculosis-specific* T helper 1 cell responses after HIV-1 infection. J Infect Dis 198, 1590–1598 (2008).

38. Amelio, P., et al. HIV Infection Functionally Impairs *Mycobacterium tuberculosis-Specific* CD4 and CD8 T-Cell Responses. J Virol 93(2019).

39. Kinter, A.L., et al. HIV replication in CD4+ T cells of HIV-infected individuals is regulated by a balance between the viral suppressive effects of endogenous beta-chemokines and the viral inductive effects of other endogenous cytokines. Proc Natl Acad Sci U S A 93, 14076–14081 (1996).

40. Casazza, J.P., et al. Autocrine production of beta-chemokines protects CMV-Specific CD4 T cells from HIV infection. PLoS Pathog 5, e1000646 (2009).

41. Buggert, M., et al. Limited immune surveillance in lymphoid tissue by cytolytic CD4+ T cells during health and HIV disease. PLoS Pathog 14, e1006973 (2018).

42. Mylvaganam, G.H., et al. Dynamics of SIV-specific CXCR5+ CD8 T cells during chronic SIV infection. Proc Natl Acad Sci U S A 114, 1976–1981 (2017).

43. Cadena, A.M., et al. Persistence of viral RNA in lymph nodes in ART-suppressed SIV/SHIV-infected Rhesus Macaques. Nat Commun 12, 1474 (2021).

44. Cecchinato, V., et al. Impairment of CCR6+ and CXCR3+ Th Cell Migration in HIV-1 Infection Is Rescued by Modulating Actin Polymerization. J Immunol 198, 184–195 (2017).

45. Perez-Patrigeon, S., et al. HIV infection impairs CCR7-dependent T-cell chemotaxis independent of CCR7 expression. AIDS 23, 1197–1207 (2009).

46. Egen, J.G., et al. Intravital imaging reveals limited antigen presentation and T cell effector function in mycobacterial granulomas. Immunity 34, 807–819 (2011).

47. Grace, P.S. & Ernst, J.D. Suboptimal Antigen Presentation Contributes to Virulence of *Mycobacterium tuberculosis* In Vivo. J Immunol 196, 357–364 (2016).

48. Yang, J.D., et al. *Mycobacterium tuberculosis-specific* CD4+ and CD8+ T cells differ in their capacity to recognize infected macrophages. PLoS Pathog 14, e1007060 (2018).

49. Muller, J., et al. Cytomegalovirus infection is a risk factor for tuberculosis disease in infants. JCI Insight 4(2019).

50. Martinez, L., et al. Cytomegalovirus acquisition in infancy and the risk of tuberculosis disease in childhood: a longitudinal birth cohort study in Cape Town, South Africa. Lancet Glob Health 9, e1740–e1749 (2021).

51. Redford, P.S., et al. Influenza A virus impairs control of *Mycobacterium tuberculosis* coinfection through a type I interferon receptor-dependent pathway. J Infect Dis 209, 270–274 (2014).

52. Beites, T., et al. Plasticity of the *Mycobacterium tuberculosis* respiratory chain and its impact on tuberculosis drug development. Nat Commun 10, 4970 (2019).

