## Supplemental Figures for "CD4 T cells are rapidly depleted from tuberculosis granulomas following acute SIV co-infection"

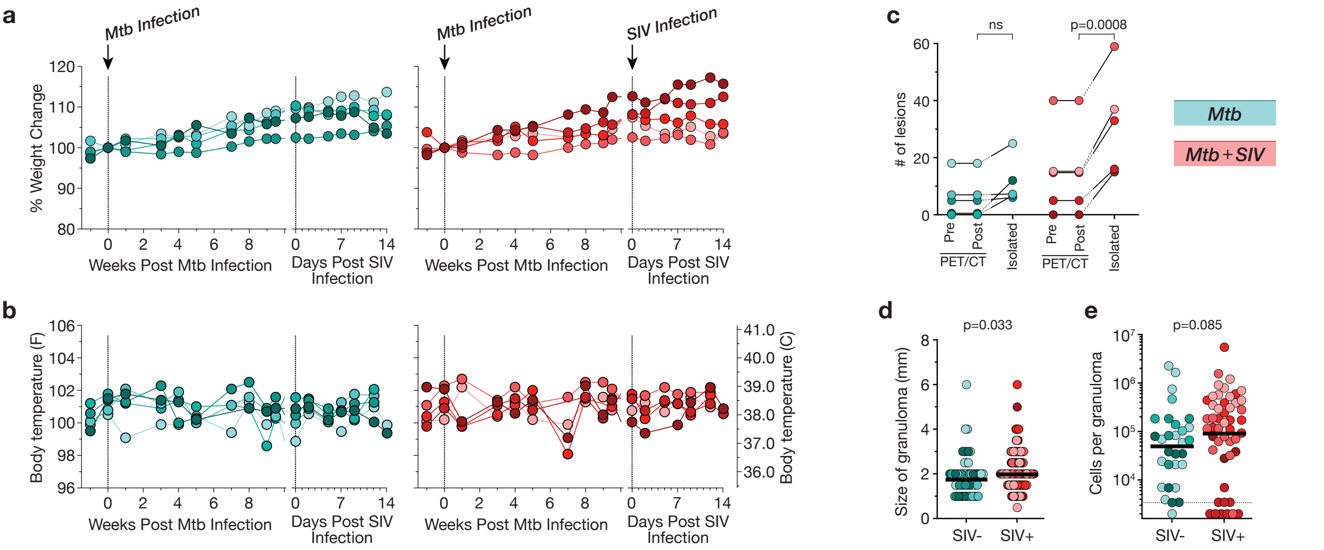


### Supplemental Figure 1

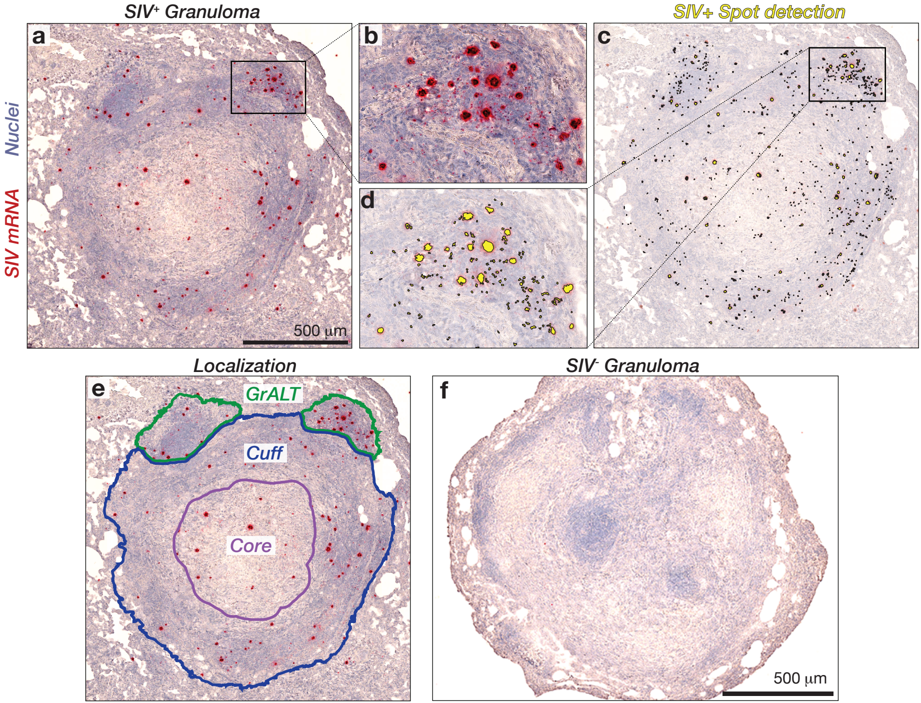


**Supplemental Figure 2**

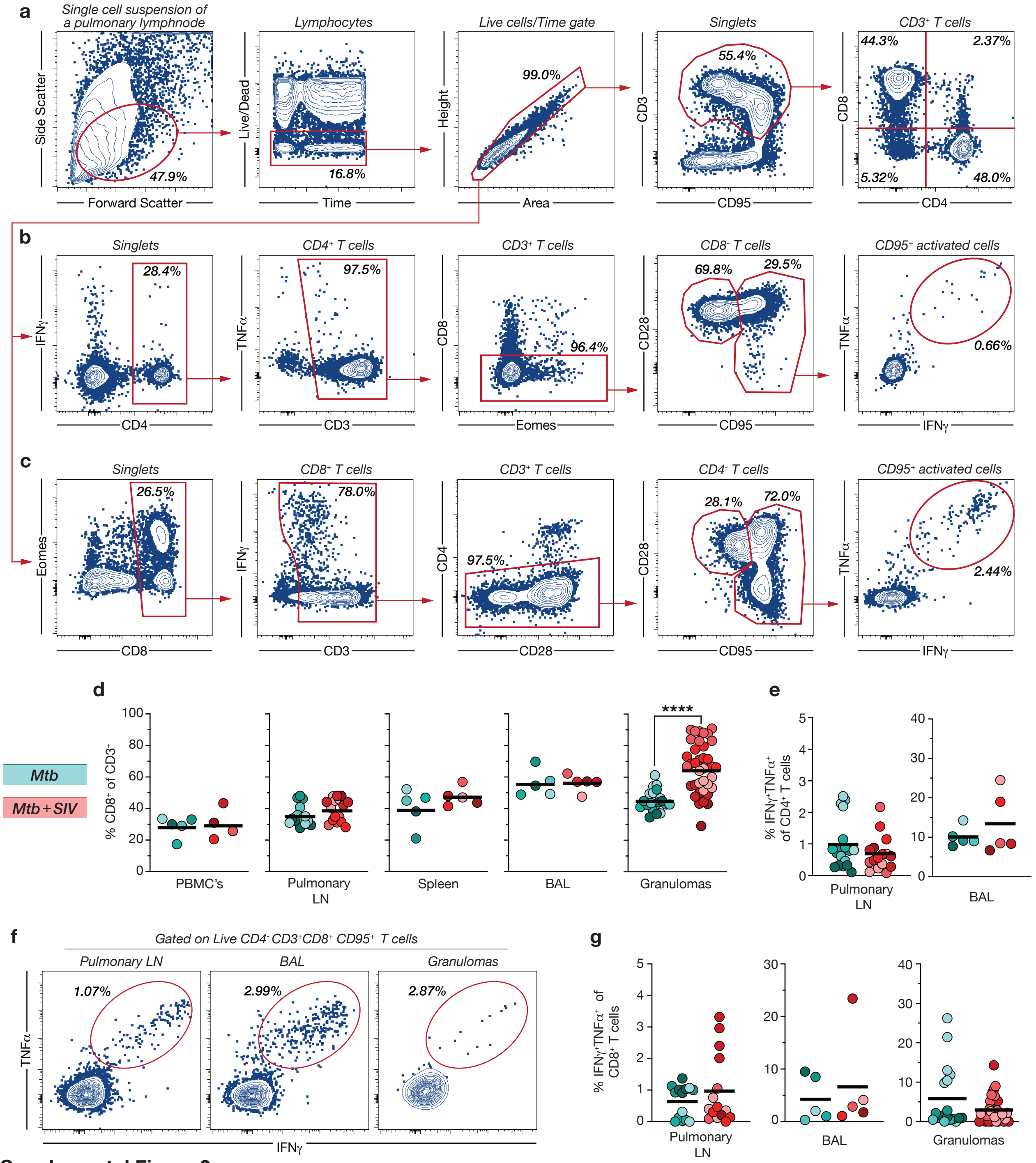


**Supplemental Figure 3**

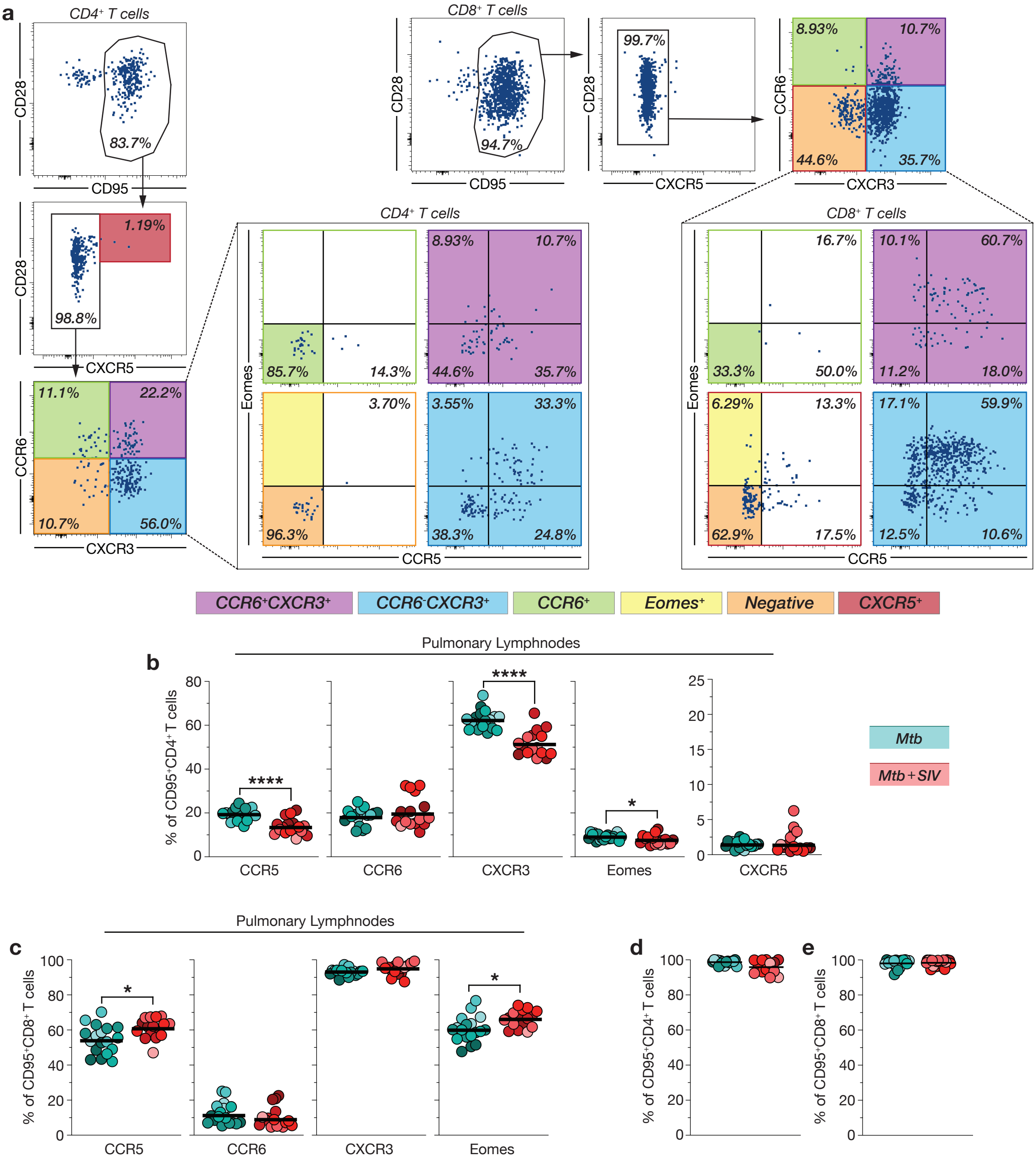


**Supplemental Figure 4**

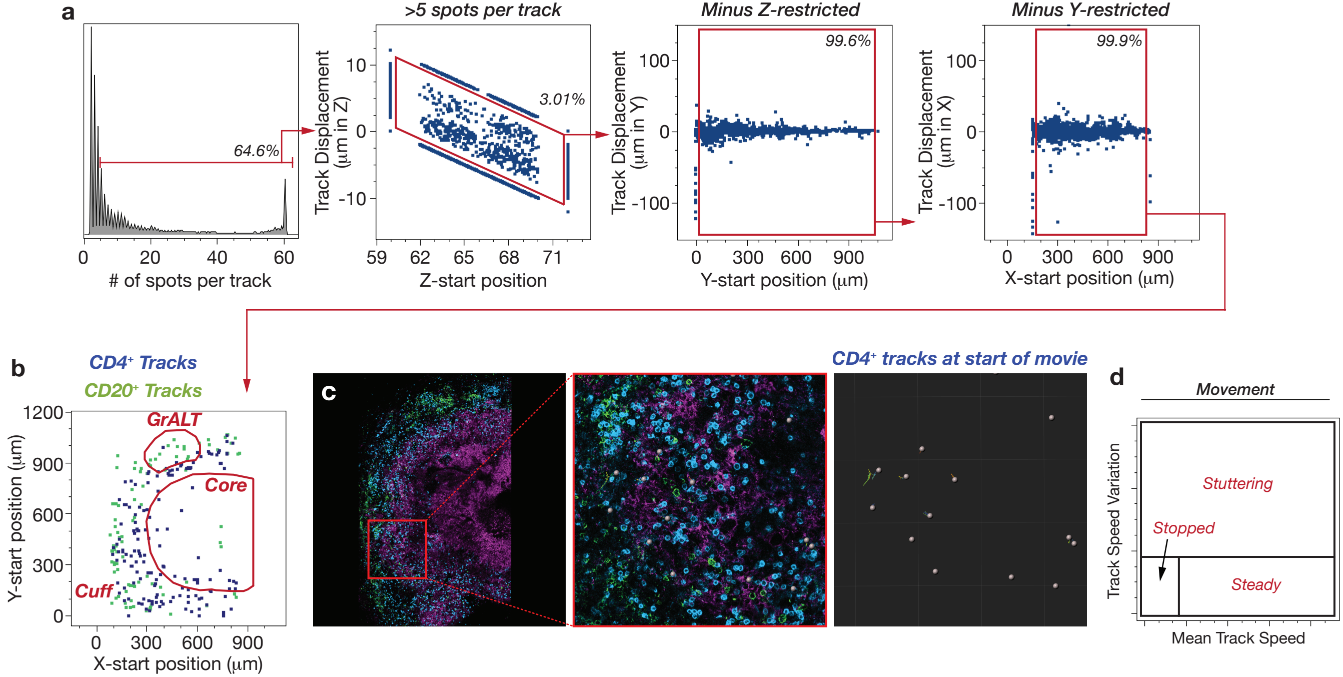


Supplemental Figure 5

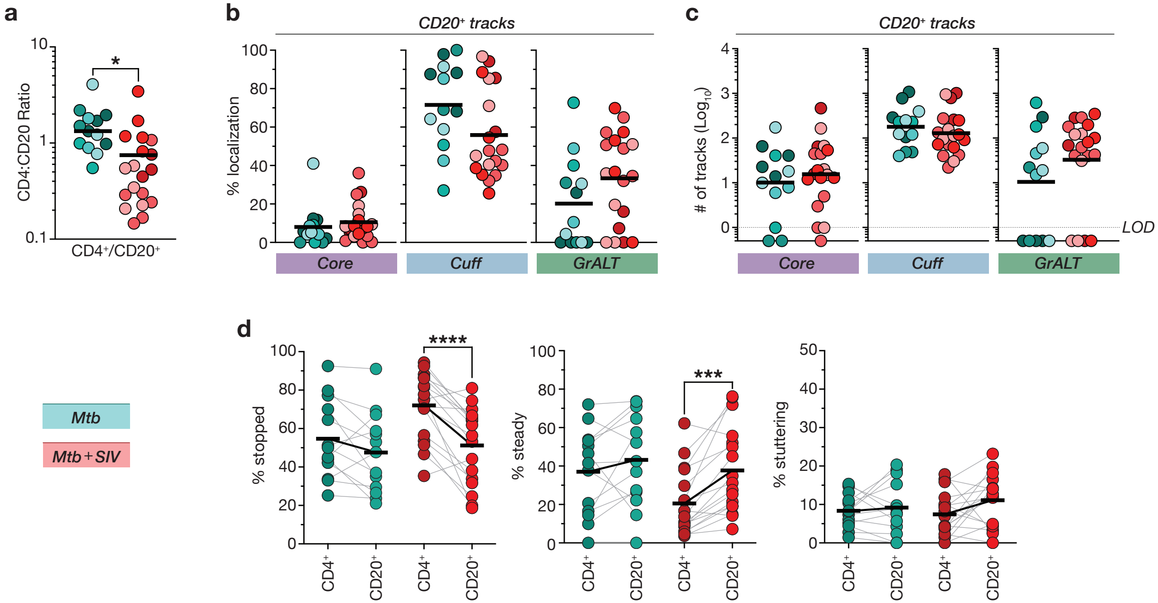


**Supplemental Figure 6**
